# MozClo: An Expanded MoClo Toolset for Large Multigene Assembly and Plant Transformations

**DOI:** 10.64898/2026.07.09.737387

**Authors:** Grant Straub, Devyn Aldrich, Cory Tobin

**Affiliations:** Mozza Foods, Los Angeles, California

**Keywords:** Golden Gate Cloning, MoClo, PhytoBricks, Plant Synthetic Biology, soybean transformation

## Abstract

The Modular Cloning (MoClo) and PhytoBrick standards have revolutionized plant synthetic biology by establishing a standardized, hierarchical assembly grammar. However, as the engineering of complex metabolic pathways, multi-trait stacks, and synthetic gene circuits expands, existing toolkits hit practical boundaries in assembly capacity and fixed grammars. To overcome these bottlenecks, we present MozClo, an expansion of the MoClo/PhytoBrick architecture. MozClo expands the standard Level 1 assembly framework to 10 positions using new L1 acceptors, end-linkers and dummy parts. We also identify and resolve a critical, sticky-end collision at L1 position 7 that has caused assembly failures during L2 cloning of large plasmids. To address commercial DNA synthesis length constraints and to lower cloning costs, we designed a universal 5-in-1 gene fragment multiplexing system. This architecture embeds up to five distinct parts flanked by orthogonal pairs of BpiI restriction sites into a single synthesized fragment, allowing them to sort independently into their respective L0 acceptor plasmids while maintaining complete modular flexibility of part types. Finally, we provide Level 2 cloning backbones with built in selection genes for common soybean transformation methods to facilitate downstream plant selection. Together, these advancements reduce DNA synthesis overhead and accelerate the construction of complex multigene payloads for plant biotechnology.

## Introduction

Plant genetic engineering has relied on the introduction of individual transgenic traits. Early successes in plant biotechnology typically relied on the expression of one or two foreign genes, frequently a single agronomic trait of interest (such as glyphosate resistance or Bacillus thuringiensis insecticidal endotoxins) paired with a plant selectable marker. Although this single-gene, single-trait paradigm successfully yielded generations of commercial genetically modified crops, it is inadequate for the next frontier of plant synthetic biology. Modern agricultural and industrial demands require plant systems to act as sophisticated, multi-functional biomanufacturing platforms. Engineering complex properties such as full metabolic pathways, gene regulatory circuits, or multi-trait resistance stacks requires the coordinated expression of numerous transcriptional units (TUs). Delivering these pathways on separate plasmids via co-transformation or attempting to combine them through generations of genetic crosses is highly inefficient; it introduces unpredictable genomic integration sites and leads to transgene segregation in successive breeding generations. Consequently, stable plant biotechnology requires the simultaneous physical delivery of large multigene payloads (often containing 5 to 10+ distinct TUs) compiled onto a single T-DNA molecule. GoldenGate assembly methods greatly simplified the challenge of combining multiple TUs in a single TDNA. Seminal toolkits, most notably the Modular Cloning (MoClo) system established by Weber et al.(1) and the parallel GoldenBraid platform(2), introduced a standardized hierarchical assembly grammar. This syntax was later codified for the wider plant research community through the PhytoBrick standard (3), which established a universal consensus for Level 0 (L0) basic genetic parts. By exploiting the ability of Type IIS enzymes to cleave outside of their recognition sequences, researchers could assemble multiple standardized basic parts into complete Level 1 (L1) transcriptional units, and subsequently stack these TUs into higher-order Level 2 constructs in efficient, single-pot reactions. Despite these elegant frameworks, scale-up constraints continue to bottleneck high-throughput plant engineering. In practice, the current standard toolkits face challenges: Transcriptional Unit Capacity: Standard MoClo architecture limits the number of transcriptional units that can be assembled in a single L2 assembly to 6 TUs. MoClo does provide for cycling through Level M or P plasmids but this adds to the time and complexity required to go from L0 parts to completed multi-gene plasmids ready for plant transformation. Sticky-End Incompatibilities: The current MoClo plasmid set contains a duplicated sticky-end in the position 7 parts which leads to very low fidelity assembly for L2 plasmids containing 7 L1 inputs. This limits the current assembly size to 6 genes. DNA Synthesis Overhead: Commercial DNA synthesis providers such as Twist Bioscience enforce minimum length thresholds. As of June 2026 Twist has a 300bp minimum for gene fragments. Ordering short regulatory elements, tags, or targeting peptides forces us to pay for filler sequences, which inflates synthesis costs. Cloning & Selection Complexity: Currently L2 multigene assemblies require that a plant selectable marker cassette be co-assembled as a separate TU. This requirement consumes a valuable cloning position. Since the number of transformation methods and selection genes are typically limited to a small number in any lab, it makes sense to include it in the L2 backbone to free up cloning positions. To address these compounding bottlenecks, we present MozClo, a new expansion of the classic MoClo/PhytoBrick standards. MozClo overcomes existing limits by expanding the L1 acceptor suite to 10 functional positions, correcting a sticky-end collision at position 7, and establishing a universal 5-in-1 gene fragment multiplexing strategy to minimize DNA synthesis cost. Finally, as a convenient utility to streamline down-stream workflows, we provide a set of Level 2 backbones including selection genes for common soybean transformation methods. Together, these advancements provide an expanded framework for multi-gene plant engineering.

## Results

### Level 1 Position 7 Sticky-End Incompatibility

We identified a f requent f ailure m ode w hen u sing t he M oClo L1 position 7 vector (pICH47791). Investigation revealed that the BpiI sticky ends derived from the pICH47791 vector overlapped with the sticky ends on L1 position 1 vector (pICH47732), resulting in incorrect or low-efficiency ligation. When assembling a L2 plasmid with 7 parts, the TGCC sticky end on the 7-part end linker (pICH50866) can ligate to the TGCC sticky end on the backbone vector, for example pAGM4673. We discovered this incompatibility after attempting to assemble 23 different plasmids that were designed with 7 TU’s. We designed the plasmids in Snapgene which indicated that the parts could in fact assemble correctly. We assembled the plasmids using standard Golden Gate reactions, cloning the TU’s into the pAGM4673 backbone. We then picked colonies, miniprepped, restriction digested the minipreps and ran the digests on agarose gels. Out of a total of 317 colonies tested, 0 had the correct band patterns, as shown in Table 1. The band patterns were mostly consistent with an empty backbone vector and no TUs inserted. We sent some minipreps to Plasmidsaurus for full plasmid sequencing and found only the end-linker part had been inserted. Our initial hypothesis was that there was a steep dropoff in assembly efficiency at 9 fragments (7 TUs, 1 backbone, 1 end linker). Since Snapgene had indicated that these parts were compatible and re-sequencing all the component parts showed all 9 fragments matched the reference files, this seemed the most likely conclusion. But repeated failures even after sampling hundreds of colonies led us to investigate the assembly by performing mock-assemblies with pen and paper. This exercise made the duplicated sticky end obvious. To resolve this problem, we redesigned the position 7 acceptor plasmid (pMOZ-L1-7F), position 7 end-linker part (pMOZ-EL-7) and position 7 dummy part (pMOZ-DUM-7) to use a CGTA sticky end in place of TGCC. We have since successfully assembled 68 L2 plasmids using the revised position 7 acceptor and end linker. The sequences can be found in the supplemental material and the plasmids available through Addgene. Table 1 includes statistics comparing the efficiency using the two different sticky ends.

**Table 1.** Frequency of successful clones determined by restriction digests. Original MoClo position 7 uses the TGCC sticky end. MozClo position 7 uses the CGTA sticky end.

|  | Correctly Assembled Clones | Total Clones Tested |
| --- | --- | --- |
| MoClo (TGCC) | 0 | 317 |
| MozClo (CGTA) | 174 | 253 |

### Expanding the MoClo Syntax to Accommodate 10 TUs

To accommodate larger genetic circuits, we expanded the standard MoClo syntax to 10 L1 positions. This expansion includes L1 acceptors, end-linkers and dummy parts. To pick new sticky ends for positions 7 through 10 we started with recommendations provided by the Lohman lab at New England BioLabs(4)(5)(6) and checked the ligation fidelity using the NEBridge Ligase Fidelity Viewer(7) and the ligation fidelity tool built in to SnapGene. This led us to choose the following pairs: GAGC-CGTA for position 7, CGTA-CTTC for position 8, CTTC-ATCC for position 9, and ATCC-ATAG for position 10. These sticky end pairs are included in the L1 acceptors and dummy parts. The end-linker parts use the following pairs to allow for linking the last L1 part back to the GGGA sticky end of the backbone: CGTA-GGGA for position 7, CTTC-GGGA for position 8, ATCC-GGGA for position 9, and ATAG-GGGA for position 10.

The full sequences for these plasmids are available in the supplemental material. The plasmids are available through Addgene. When using these plasmids in 12 plasmid assemblies (10 TUs, 1 end-linker, 1 backbone) some adjustments must be made to the GoldenGate reactions recipe to accommodate this many inputs while also keeping the volume of each miniprep reasonable for pipetting.

Here is a suggested recipe for a 12 part assembly: 0.5uL backbone miniprep, 1.0uL each of 10 Level 1 minipreps, 1.0uL T4 ligase, 1.0uL BpiI, 1.4uL 10X T4 ligase buffer. As more parts are added we keep the volume of enzymes added the same but increase the buffer so the final proportion of the buffer is 10% of the entire reaction volume.

### Maximizing DNA Synthesis via Gene Fragment Multiplexing

Twist Bioscience imposes a 300bp minimum for gene fragments. To prevent paying for empty padding sequence when ordering short parts, we built a collection of universal L0 acceptor plasmids that allows one to multiplex from 1 to 5 parts on a single gene fragment. Each part is flanked by a unique, orthogonal pair of BpiI sticky ends. The Eco31I sites that define the part type are located inside of the BpiI sites.

Following the original MoClo standard, gene fragments would have a single BpiI site added to each end and the Eco31I sites would be added through a shared sticky end on the L0 acceptor plasmid. This limits the possible part types to those defined by the L0 acceptor plasmid collection. Our company needed more flexibility to accommodate, for example, parts that spanned NTAG-CDS2 or custom sequences upstream of the start codon to control the Kozak sequence. Table 3 shows recommended Eco31I sites for each part type. But since practically any sticky end can be synthesized, one is not limited to these sticky ends. For example, we frequently use ATGA at the junction between 5UTR and NTAG parts. This forces the first nucleotide after the start codon to be A.

**Fig. 1.**
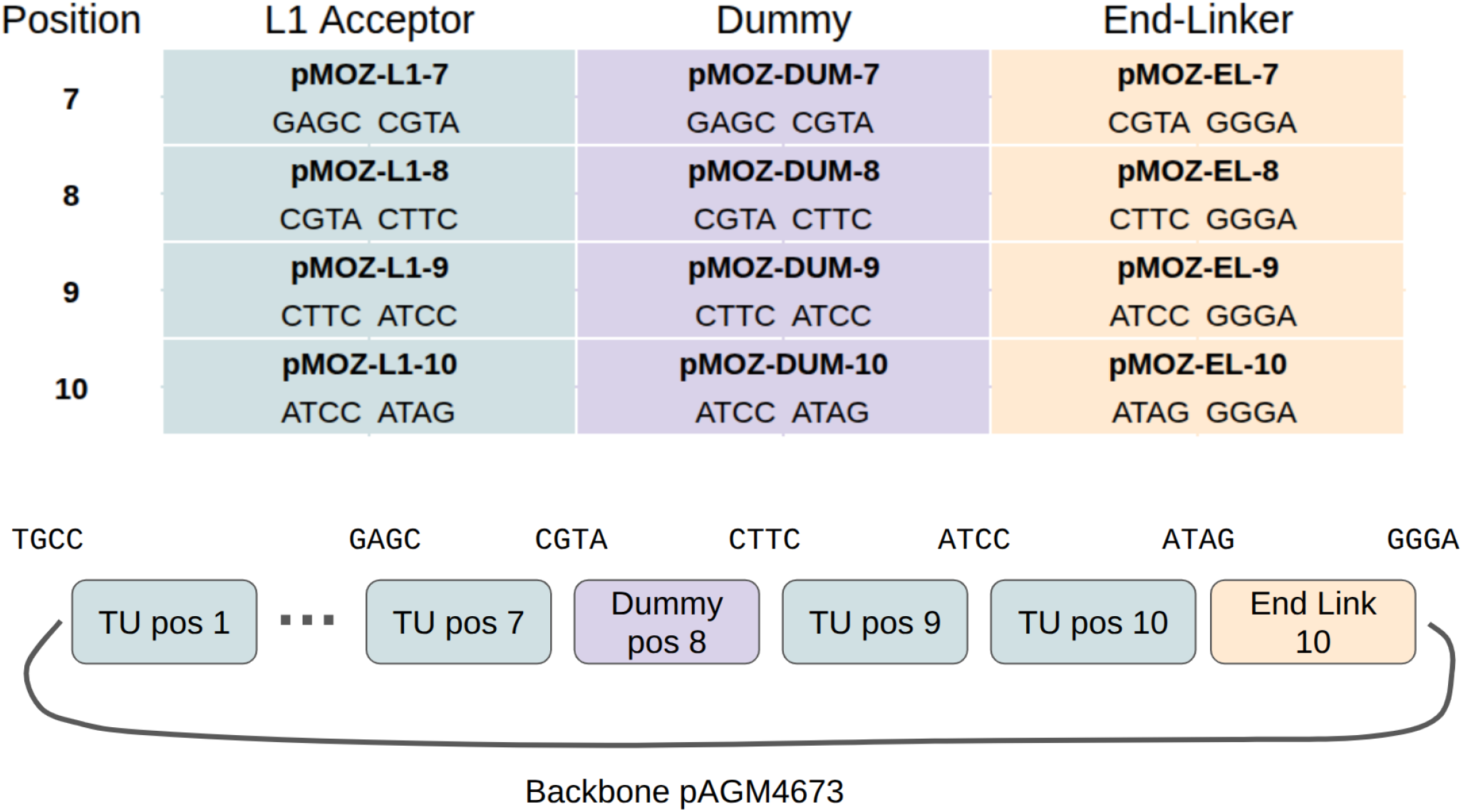
Top: Sticky ends used for expanding the MoClo syntax to 10 positions and corresponding plasmid names. Bottom: Assembly of 10 TUs, end linker, and backbone into a finished L2 plasmid.

But allows for us to to completely define the upstream Kozak sequence on the 3’ end of the 5UTR part during synthesis. Additionally, parts can be made that span these definitions. For example, we frequently design coding sequences that span the NTAG to CDS2 parts because we do not need any modularity in the majority of the coding sequence but do want to be able to swap epitope tags or targeting peptides on the C-terminus. A part designed for this use case would have the GGTCTCT**CCAT** adapter on the left (5’) side and the **TTCG**AGAGACC adapter on the right (3’) side.

To design a gene fragment utilizing the universal L0 acceptors, start with the sequence of the genetic part. Add Eco31I sites to the 5’ and 3’ ends of the part, making sure to get the reading frame correct in scenarios where that is important, such as the CDS1 left adapter. Then pick 1 of the 5 BpiI adapters from Table 2. Add the left adapter sequence to the 5’ end of the part, outside of the Eco31I site. Then add the right adapter sequence to the 3’ end of the part. Do this for each of the parts to be included in a single gene fragment. Merge up to 5 parts together in any order, making sure (1) to not duplicate the BpiI site, and (2) add 8bp of padding between adjacent BpiI sites. If the total length of the combined sequence does not add to 300bp, add random filler sequence to ends. An example of a 5-part gene fragment is included in the supplement.

**Table 2.** Column 2 shows the BpiI sticky ends to be used when designing multiplexed gene fragments. Column 3 and 4 show the BpiI adapter sequences to be added to the left and right side of each part within the gene fragment. Including the BpiI recognition sites.

| Universal L0 Acceptor | Flanking BpiI Overhangs | Left BpiI Adapter | Right BpiI Adapter |
| --- | --- | --- | --- |
| pMOZ-L0-A | CTCA / CGAG | GAAGACAACTCA | CGAGTGGTCTTC |
| pMOZ-L0-B | GGAG / TACT | GAAGACAAGGAG | TACTTGGTCTTC |
| pMOZ-L0-C | CCAT / AATG | GAAGACAACCAT | AATGTGGTCTTC |
| pMOZ-L0-D | AGGT / TTCG | GAAGACAAAGGT | TTCGTGGTCTTC |
| pMOZ-L0-E | GCTT / CGCT | GAAGACAAGCTT | CGCTTGGTCTTC |

**Table 3.** Eco31I adapter sequences to include in the gene fragment, inside of the flanking BpiI adapter sequences.

| Type | Left Adapter | Right Adapter |
| --- | --- | --- |
| Pro | GGTCTCTGGAG | TACTAGAGACC |
| 5UTR | GGTCTCTTACT | CCATAGAGACC |
| NTAG | GGTCTCTCCAT | AATGAGAGACC |
| CDS1 | GGTCTCTAATG | AGGTAGAGACC |
| CDS2 | GGTCTCTAGGT | TTCGAGAGACC |
| CTAG | GGTCTCTTTCG | GCTTAGAGACC |
| 3UTR-TERM | GGTCTCTGCTT | CGCTAGAGACC |

### L2 Backbones for Plant Selection

The original MoClo specification does not provide plant selection directly in the Level 2 cloning backbones, aside from pICH86966 which provides NOS::Kan selection. Generally, users of the MoClo kit would build a L1 plasmid with the selection gene of their choice.

This would get included in the final plasmid as one of the TUs in the last BpiI GoldenGate reaction. This provides modularity but also consumes one of the L1 cloning positions. We found that we always used the same selection gene for transformation so the modularity provided was not needed. But the additional slot for an extra TU would be useful. Therefore we designed a set of L2 acceptors with plant selection genes inside the TDNA boundaries but outside of the cloning site. See Table 4

**Table 4.** Level 2 plasmids containing genes for in-plant selection.

| Plasmid | Plant Selection | Selection Gene |
| --- | --- | --- |
| pMOZ-L2-AHAS | Imazapyr | StUBI3::AHAS::StUBI3 |
| pMOZ-L2-Hyg | Hygromycin | StUBI3::HygR::StUBI3 |
| pMOZ-L2-Bar | Glufosinate | NOS::Bar::NOS |

Recently there has been progress on increasing the efficiency and throughput of soybean transformations. But these new methods do not use the same selection agents that are common in Arabidopsis (Basta and kanamycin). Embryonic axis transformation methods(8)(9)(10) use a mutated version of the AHAS gene that is resistant to the herbicide Imazapyr. We built a plasmid pMOZ-L2-AHAS that contains the UBI3 promoter and terminator from potato to drive the expression of the AHAS gene. The AHAS protein has a serine to asparagine mutation at position 653.

Cotyledon node transformation methods such as the one described by Luth et al(11), rely on glufosinate (Basta) selection. We built a plasmid pMOZ-L2-Bar that uses the NOS promoter and terminator to drive expression of the Bar protein. Finally, soybean somatic embryo transformations(12) and soybean in-planta transformation methods(13) both use hygromycin selection. We built the plasmid pMOZ-L2-Hyg to accommodate these methods. It utilizes the potato UBI3 promoter and terminator to drive expression of the aminoglycoside phosphotransferase protein (HygR) from Ecoli to provide hygromycin resistance.

**Fig. 2.**
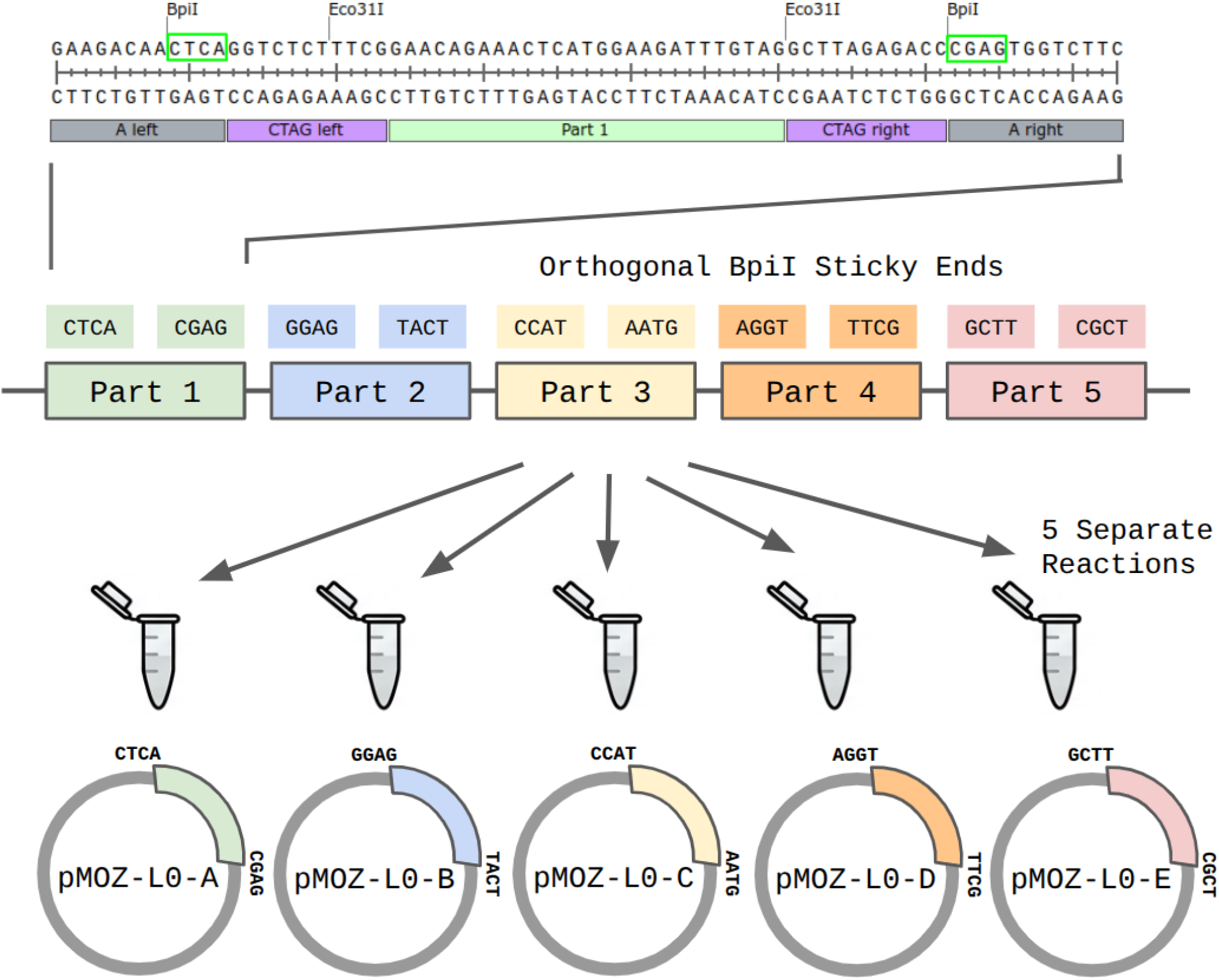
Top: Example sequence of a single part flanked by Eco3I sites, further flanked by BpiI sites. Bottom: Schematic of the orthogonal pairs of BpiI sites directing fragments from a single molecule into 1 of 5 L0 acceptor plasmids.

Additional design considerations were made with these plasmids beyond plant selection genes. The original MoClo L2 acceptors used multiple genes to produce the orange pigment β-carotene to aid in identifying colonies with undigested backbone. In our laboratory conditions, the development of the orange pigment sometimes added an additional 12 hours to the growth time before we could reliably pick non-orange colonies. So we designed these plasmids with lacZ for blue white screening which develops faster. Although, this does add additional cost for the media which needs IPTG and X-Gal. We also included a noncoding, 256bp sequence called “tagurit” that we use as a universal target for qPCR assays to determine insert copy number or zygosity. All our TDNA’s include this sequence so qPCR assays can be performed identically regardless of the other genes in the TDNA. And since it is adjacent to the right boundary, we also use tagurit as an oligo binding site in insert mapping assays.

## Discussion

By resolving the sticky end overlap between MoClo positions 7 and 1, we fixed an assembly problem that had long plagued our lab. We hope this bug fix is useful for the rest of the plant synthetic biology community. Additionally, expanding the number of level 1 TU’s to 10 allowed us to build plasmids that incorporated multiple recombinant proteins, chaperones, regulatory proteins and regulatory RNAs all in a single TDNA. We believe that the future of plant synthetic biology will require more complex gene circuits than has been previously possible and that the MoClo cloning system will need to be expanded further. Research coming out of the Lohman lab at New England Biolabs provides a path forward to at least 18 TUs on a single TDNA. It is to be determined if the increased length of the TDNA will be accommodated by current plant transformation methods.

The impetus for the development of multiplexed gene fragments was to save money on synthesis when frequently buying small (<300bp) parts. Later we realized that by building the Eco31I sites into the gene fragment, we had greater flexibility in designing non-standard parts since we are not tied to specific part-type-specific L0 acceptors. This has allowed us to better control which parts of our genes we chose to break up into modular sections. It has also allowed us to explore certain sequences which would have been prohibited by the original MoClo syntax. For example, we have seen recombinant protein expression increases by controlling the sequences immediately upstream of the start codon.

Currently the merging of parts onto single gene fragments is done manually in Snapgene which sometimes leads to less optimal layouts that miss some potential financial savings. To fully realize the potential of the MozClo multiplexing system at scale, there is a need for the development of dedicated computer-aided design software tools to assist in the gene fragment layout. An ideal software solution would take a part with Eco31I sites already in place. It would then arrange the parts together on different gene fragments, using the Twist Bioscience API to assess manufacturability while also maximizing sequence-length utilization and then add appropriate BpiI adapters to each part.

Developing such open-access software tools will be essential to lower the barrier of entry for MozClo, transitioning the design of highly multiplexed gene fragments from a tedious manual chore to a streamlined, automated press-of-a-button pipeline. This integration of wet-lab innovation and software automation represents the natural trajectory of synthetic biology, making complex plant genetic engineering faster, more reliable, and universally accessible.

Arguments for multiplexing are that it allows for greater fleibility in part design and saves money on groups of small parts. An argument against multiplexing is that it adds another layer of complexity to designing genetic components. Nevertheless, we hope this is beneficial for advancing plant synthetic biology.

## Materials and Methods

### Plasmid Construction and Architecture

The MozClo plasmid library was generated using standard GoldenGate cloning methods starting with Gene Fragments from Twist Biosciences. The exact methods are irrelevant but the Snap-gene files for each plasmid are available in the supplemental material and contain the full assembly scheme in the Snap-gene history. The source material for building these plasmids was primarily synthetic DNA and plasmids from the MoClo kit (Addgene kit#1000000044) which was a gift from the Sylvestre Marillonnet lab.

### Measuring the Rate of Successful Assemblies

To assess the efficiency of using CGTA instead of TGCC sticky ends on position parts, we designed many different plasmids that utilized those sticky ends. We assembled them with the following recipe: 0.5uL backbone miniprep, 1.0uL miniprep of each of 7 level 1 minipreps, 1.0uL of the end-linker miniprep, 1.2uL ThermoFisher T4 10X ligase buffer, 1.0uL ThermoFisher FastDigest BpiI, 0.5uL nuclease free water. In a thermalcycler, the temperature was cycled between 37°C for 2.5 minutes and 16°C for 3 minutes, 34 times. Followed by a single step of 50°C for 5 minutes and then 80°C for 10 minutes. The reaction was then electroporated into Lucigen E. cloni 10G competent cells with a BioRad MicroPulser, 0.2cm cuvettes and the EC2 program. 110uL of LB was added and incubated at 37°C for 45 minutes. The liquid was then plated on LB plates containing 50ug/mL kanamycin, 24ug/mL IPTG and 33ug/mL X-Gal (*note for X-Gal we use DMSO as the solvent). Multiple colonies for each plate were picked and grown in 5mL LB with kanamycin. They were each miniprepped with the Macherey-Nagel Nucleospin Plasmid Miniprep kit. For each design, appropriate FastDigest restriction enzymes were chosen and reacted with the minipreps according to the FastDigest product manual, for 90 minutes at 37°C. The completed reactions were loaded into 1% agarose TAE gels and run alongside a 1kb plus DNA ladder. The image of the gel was compared against a simulation in Snapgene. If the bands in the image match the simulation, it was counted as a successful assembly, even if later full plasmid sequencing showed a point mutation somewhere in the plasmid. For each plasmid with at least one correct band pattern, one miniprep was sent for full plasmid sequencing with Plasmidsaurus. The results of this were not counted as part of the success/failure ratio. A small quantity of minipreps with incorrect band patterns were sent for full plasmid sequencing for debugging purposes. Please note we did not design this as an experiment to test the efficiency of different sticky ends on position 7 plasmids. These plasmids were designed and built as part of our company’s plasmid assembly workflow. This experiment was reconstructed later using the data we collected and saved as part of that workflow.

### Plasmid Availability

All plasmids described in this manuscript have been deposited to Addgene accessible from the URL https://www.addgene.org/Cory_Tobin/

## Supporting information

Assembly stats by plasmid

Example sequences

Plasmid Sequences

## ACKNOWLEDGEMENTS

Thank you to the Marillonnet, Patron, Tissier, and Orzaez labs for pioneering the molecular methods that have made plant synthetic biology possible and publishing them open access.

## Notes

https://www.addgene.org/Cory_Tobin/

### Competing Interest Statement

The authors have declared no competing interest.

